# Predator diversity and thermal niche complementarity attenuate indirect effects of warming on prey survival

**DOI:** 10.1101/2020.09.29.319392

**Authors:** Adam Pepi, Marshall McMunn

## Abstract

Climate warming has broad-reaching effects on communities, and although much research has focused on direct abiotic effects, indirect effects of warming meditated through biotic interactions can be of equal or greater magnitude. A body of theoretical and empirical work has developed examining the effects of climate warming on predator-prey interactions, but most studies have focused on single predator and prey species. We develop a model with multiple predator species using simulated and measured predator thermal niches from a community of ants, to examine the influence of predator diversity and other aspects of community thermal niche on the indirect effects of climate warming on prey survival probability. We find that predator diversity attenuates the indirect effect of climate warming on prey survival probability, and that sufficient variation of predator thermal optima, closer prey and mean predator thermal optima, and higher predator niche complementarity increases the attenuation effect of predator diversity. We predict therefore that more diverse and complementary communities are likely more affected by direct versus indirect effects of climate warming, and vice versa for less diverse and complementary communities. If general, these predictions could lessen the difficulty of predicting the effects of climate warming on a focal species of interest.

## Introduction

Understanding the effects of climate drivers on population dynamics has become an important goal in ecological research, with the ultimate aim of predicting the effects of climate change on communities and ecosystems (Walther et al. 2002; Parmesan 2006). A main driver of climate effects on population dynamics is warming, which affects ectotherms primarily through increased metabolic rate (Uszko et al. 2017). The direct effects of increased metabolic rates on ectotherms are complex and depend on details of life history, but some predictions are possible based on sufficient life history information (e.g., Bale et al. 2002; Deutsch et al. 2018). However, the effects of climate change on ecological communities is complicated by biotic interactions, and many effects of climate change on population and community dynamics are likely to be indirect and mediated through biotic interactions (Tylianakis et al. 2008; Post 2013), which cannot be predicted based solely upon the life-history of the focal species. This complexity makes predicting the impacts of climate warming on individual species, let alone whole communities, extremely difficult.

Recognizing the importance of biotic interactions as mediators of climate change effects, some research has investigated which factors influence climate’s effects on interactions. For example, mechanistic models and experimental work suggest that the equilibrium population size of predators and prey with warming can be predicted based on predator foraging mode and asymmetries in the relative response rates of predator and prey movement velocity to temperature (Dell et al. 2014; Ohlund et al. 2014). Other work suggests that asymmetries in relative response rates of predator attack rate and development rates of prey with size- or stage-dependent refuges will also affect equilibrium population sizes (Culler et al. 2015; Pepi et al. 2018).

Thus far, most research has focused on single predator-prey pairs. However, most prey species likely interact with multiple predators, all of which will be affected by a warming climate in ways that vary with species’ life histories (Barton and Schmitz 2009). Predator diversity is likely to have important impacts on how climate warming affects prey species population size, due to the compensatory effects on many trophic links. These compensatory effects might arise due to variation in predator thermal niches that are complementary with respect to activity or fitness in a given range of temperatures (*i.e.,* low thermal niche overlap). In this situation, the change in metabolic rate and thus attack rate in response to temperature by one predator might be concurrent with changes in another predator that may compensate for the change in attack rate by the first. Alternatively, variation in predator thermal niches may not result in compensation if thermal niches are similar (*i.e*., a large overlap): with very similar thermal niches, effects of warming on predation might be even more intense than expected for a single predator-prey pair, as attack rates for both predators increase or decrease together with temperature.

How warming will affect predator-prey interactions considering multiple predators likely depends on the community distribution of thermal response curves, or the community thermal niche (Kühsel and Blüthgen 2015). However, there has been very limited empirical work in this area. Kühsel and Blüthgen (2015) examined flower visitation rates by pollinators in agricultural meadows, and characterized the thermal niche of 511 species of arthropods based on activity rates across a range of temperatures, finding that higher thermal niche complementarity within communities resulted in greater predicted resilience of pollination function with warming. Systematic variation with insect order was observed, with a great degree of variability of thermal niches across species within each community (s.d. of thermal optimum from 3.1-5.9°C in different taxa). There was also variation between communities in thermal niche complementarity, which was defined as a weighted coefficient of variation of thermal optima within the community.

For a community with substantial thermal niche variation, if prey species are consumed by multiple predators, it is likely that there will be both positive and negative asymmetries in thermal response rates between the prey species and predators. From this perspective, we can predict that temperature response of a given prey species relative to all the temperature responses of the predators with which it interacts will determine whether climate effects mediated by predation lead to an increase or decline of the prey species. This means that for prey populations interacting with a single or few predator species it is very likely that asymmetries will be an important mediator of climate effects of population levels. However, as the number of interacting predator species increases, the likelihood that a large net asymmetry in temperature responses between predator and prey will remain becomes lower, if the predator thermal niches are not all warmer or cooler than the prey. In this latter situation, we would expect that the indirect climate effects mediated by biotic interactors become less important, and thus the direct effects of warming should be more important.

To investigate the effects of predator diversity on prey abundance due to thermal rate asymmetries in a warming environment, we developed a simulation model of predator-prey communities. For this study, we use the example of a univoltine invertebrate prey species with a temperature-dependent window of vulnerability, based on data from an Arctiine moth (*Arctia virginalis*; Pepi et al. 2018). This type of life-history, in which a species is vulnerable to predators during a juvenile stage or below a certain size threshold that is approached at a temperature-dependent growth rate, is common across taxa (e.g., insects: Benrey and Denno 1997; amphibians: Werner 1986; marine organisms: Paine 1976, Mittelbach 1981, Christensen 1996). We used simulated communities of predators to test our prediction that increased predator diversity will weaken indirect effects of warming. We also characterized the thermal niche of a community of 21 species of ants in California as an empirical example of our simulated predator communities. We conducted sensitivity analyses to examine under which conditions we would expect predator diversity to attenuate the indirect effects of warming. We conducted sensitivity analyses on the range of variation (s.d.) of predator thermal optima, predator thermal niche breadth, and the relative value of the predator community mean thermal optimum versus the prey species thermal optimum on changes to prey survival probability with warming.

## Methods

### Simulation model

We developed a demographic model to simulate temperature-dependent survival probability across one generation of a single prey species, interacting with one or many predator species, extending a model previously developed in Pepi et al. (2018). This model combines temperature-dependent growth of prey, size-dependent predation on prey, and temperature-dependent attack of predators to simulate predation risk *P* at temperature *T*:

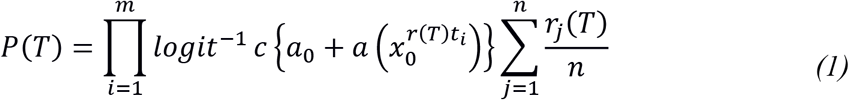

in which *t*_*i*_ is time in days, *m* is the length of the simulation in days, *x*_0_ is log initial caterpillar mass (10 mg), *T* is temperature in Celsius (from 10-30 °C), *r* is instantaneous daily growth rate of the prey as a function of temperature, *a*_0_ and *a* represent the intercept and slope of size-dependent daily predation, *c* is a constant to scale size-dependent predation at the starting size to 1, *n* is the number of predators in the community, *r*_*j*_ is the daily predation rate of the predator *j* on prey as a function of temperature. To determine predation risk at each temperature, daily size-dependent predation risk is calculated based on prey size (scaled relative to a maximum predation rate of 1 at *x*_0_), which in turn is calculated based upon temperature-dependent exponential growth. Size-dependent predation risk is multiplied by the average attack rate of all predators in the community. The temperature-dependent growth rate (*r*) of prey and attack rate of predators are modelled as a unimodal Gaussian function of temperature (*T*) after Taylor (1981):

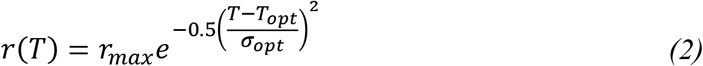

in which *r*_*max*_ is the maximum rate, *T*_*opt*_ is the thermal optimum, and *σ*_*opt*_ determines the spread of the unimodal curve. For predators, the thermal optimum is drawn from normal distribution with mean *μ* and s.d. *σ*^2^.

Simulations were conducted of predator communities including 1, 3, 10, and 30 predators to assess the effects of predator diversity on prey under warming. Size-dependent predation risk was parameterized based on field data from *Arctia virginalis* caterpillars preyed upon by *Formica lasioides* (Pepi et al. 2018; Table 1). Temperature-dependent survival and growth of *A. virginalis* caterpillars reared in incubators from 10-25° C were estimated using equation 2 and used to compare the direct vs. indirect effects of warming (Pepi et al. 2018). A mean thermal optimum of 23.5 °C (to match the thermal optimum of *A. virginalis*; Table 1) was used for the distribution of the predator community, with an s.d. of 5, which is within the upper range of variation documented by Kühsel and Blüthgen (2015) in natural arthropod communities.

**Table 1.** Values of parameters used for simulations, or values when parameters were held constant in sensitivity analyses.

| Parameter | Description | Value when constant | Source |
| --- | --- | --- | --- |
| Size-dependent predation: |  |  | Pepi et al 2018 |
| $a_0$ | Intercept | 2.7576 | |
| $a$ | Slope | -1.438 | |
| $x_0$ | Starting size | 10 mg | |
| $c$ | Scaling constant | 2.739361 | |
| Prey growth: |  |  |  |
| $r_{max}$ | Maximum rate | 5.9 | Fitted from rearing data in Pepi et al. 2018. |
| $T_{opt}$ | Thermal optimum | 23.5°C | |
| $\sigma_{opt}$ | Niche breadth | 6.7°C | |
| Prey rearing survival: |  |  | Estimated from unpublished data: rearing method in Pepi et al. 2018. |
| $r_{max}$ | Maximum rate | 0.75299 | |
| $T_{opt}$ | Thermal optimum | 17.661°C | |
| $\sigma_{opt}$ | Niche breadth | 7.9°C | |
| Predator attack rate: |  |  |  |
| $r_{max}$ | Maximum rate | 0.8 | $r_{max}$ extrapolated from data in Karban et al. 2015, $\mu$ of $T_{opt}$ set equal to $T_{opt}$ of prey, $\sigma$ of $T_{opt}$ and $\sigma_{opt}$ chosen based on values in Kühsel and Blüthgen 2015. |
| $\mu$ of $T_{opt}$ | Mean of thermal optimum | 23.5°C | |
| $\sigma$ of $T_{opt}$ | Standard deviation of thermal optimum | 5°C | |
| $\sigma_{opt}$ | Niche breadth | 4°C | |

For each simulation, the indirect effect of warming on survival (Δ_*indirect*_) was calculated. We define this as the net change (positive and negative) in the difference between rearing survival and survival including rearing survival and predation risk with respect to temperature *T* between minimum and maximum *T*. We calculate this value by integrating the absolute value of the derivative with respect to *T*:

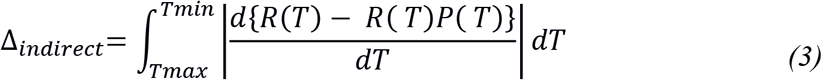

in which *R*(*T*) is rearing survival at temperature *T*, and *P*(*T*) is predation risk at temperature *T* as defined in equation 1. We used the absolute value derivative because we are interested in how much net change is due to the indirect action of warming, in a positive or negative direction.

According to the net change theorem and the definition of an absolute value, equation 3 simplifies to:

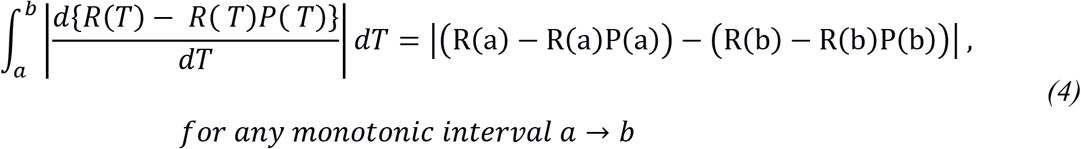

To identify sub-intervals that are monotonic we fit a third order polynomial function fit in the following form:

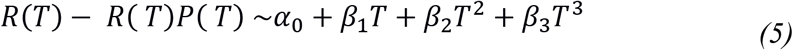

We then calculated all real roots of the derivative of this polynomial (5) to define monotonic sub-intervals. We calculated the absolute value of change within these sub-intervals as in equation 4, and summed all intervals for net change between minimum and maximum temperature. For each predator community size, 10,000 simulations were conducted and the distribution of Δ_*indirect*_ was summarized.

To test the broader applicability of the simulation model beyond the particular parameter values chosen for the initial constant parameter model, and to understand the broader ecological implications of the results, we conducted univariate sensitivity analyses. Model sensitivity to community thermal niche parameters was analysed by individually varying the s.d. and mean of the distribution of predator thermal optima, and the spread of predator thermal niches (*σ*_*opt*_).

### Empirical data: ant community thermal niche

Thermal niches of ants were estimated by pairing passively collected foraging ants with detailed ground surface temperature measurements. Collections were made using automated time-sorting pitfall traps, which simultaneously measure ground-surface temperature and separate active ants into 24 separate hour-long pitfall samples (McMunn 2017). To install traps, we carefully removed leaf litter and dug a small hole approximately 20 cm wide, 30 cm long, and 20 cm deep. We then buried the trap; replacing soil flush with the top the trap entrance, a funnel coated with fluon (Insect-a-slip, Bioquip, Rancho Cordova, CA). We replaced leaf litter, taking care to minimize the disturbance to the surrounding litter and soil. After installation, the traps remained closed to ants for 24 hours to avoid ants being initially attracted to the soil disturbance following pitfall trap installation (Greenslade 1973). We separated ants from all other collected arthropods, identified each individual to species, and after confirmation of species identifications by Phil Ward (UC Davis), deposited vouchers at the Bohart Museum of Entomology (UC Davis).

The study area consisted of 4.2 hectares of mixed sagebrush shrubs and coniferous forest in the Sierra Nevada mountain range (2000 m, 39.435583°N, 120.264017° W). The range of *A. virginalis*, which was used to parameterize the prey thermal response in the simulation model, extends to this geographic area. We made collections between 19 June and 14 October 2015, concentrated in one- or two-week sampling bouts each month. From a grid of 600 potential sites across the habitat, we randomly selected 127 sites for 24 hourly ant collections. Over the season, this resulted in 3048 hour-long samples of ant abundance (24 hrs × 127 sites = 3048 hourly samples). The traps recorded temperature measurements every 5 minutes during collections using a K-type thermocouple datalogger at a height of 1-3 mm above the surface of the leaf litter or above the surface of the soil if no litter was present (McMunn 2017).

Ant thermal niche was modelled by species with equation (2) fit using nls (package nlme, Pinheiro et al. 2017), using abundance data aggregated across all sampling locations and dates. Fitted parameter values for 20 ant species that were sufficiently abundant to successfully fit thermal response curves were used to generate simulated communities of varying diversity. Species were drawn randomly from this set of 20 without replacement for simulations, using the same procedure as described above, except that values of *T*_*opt*_ and *σ*_*opt*_ were fitted values from the ant community data. Since there were large differences in abundance between ant species in this community (which is dominated by 2-3 species), simulations were conducted both with estimated *r*_*max*_ values (maximum activity rate by species), and with all values of *r*_*max*_ = 0.8, to create an even distribution of species.

## Results

Individual simulations generally showed more stability in overall predation rate over the temperature range with higher predator diversity (Figure 1), and the results of many simulations together showed a decrease in Δ_*indirect*_ with higher predator diversity (Figure 2a) with the initial constant parameter values (Table 1). In simulations using empirical ant community data, greater ant diversity did not generate a decrease in Δ_*indirect*_ with fitted *r*_*max*_ values (Figure 2b), but with all values of *r*_*max*_ = 0.8, greater ant diversity did lead to a decrease in Δ_*indirect*_(Figure 2c).

**Figure 1.**
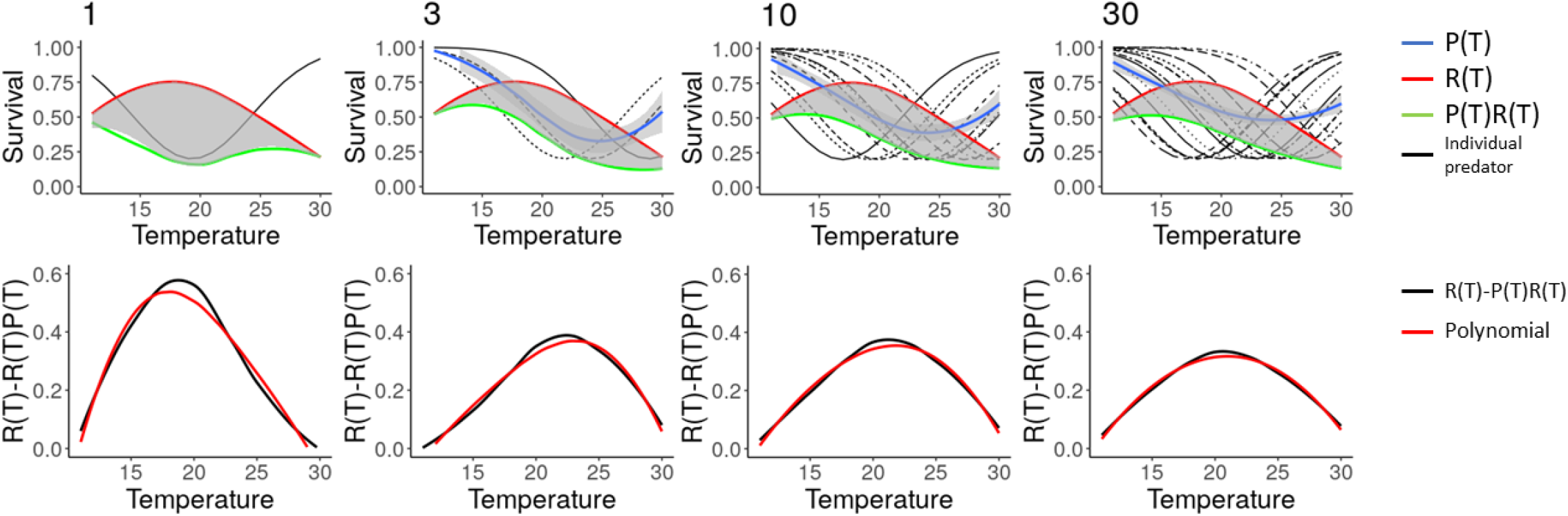
Plot displaying simulated prey survival with increasing temperature and indirect effects of warming on prey [R(T)-R(T)P(T)] preyed upon by 1-30 predator species, from left to right. Black lines show (1 - predation rate) of each predator, the blue line shows (1- mean predation rate) of all predators with a 95% CI, the red line shows rearing survival of prey [R(T)], the green line shows the combined effects of predation and rearing on survival [R(T)P(T)], and the grey band shows the indirect effect of warming on survival [R(T)-R(T)P(T)]. The indirect effect of warming [R(T)-R(T)P(T)] is plotted in the second row, with the black line showing R(T)-R(T)P(T) (the grey band in the first row), and the red line showing the fitted cubic polynomial. This plot displays representative results of a single simulation.

**Figure 2.**
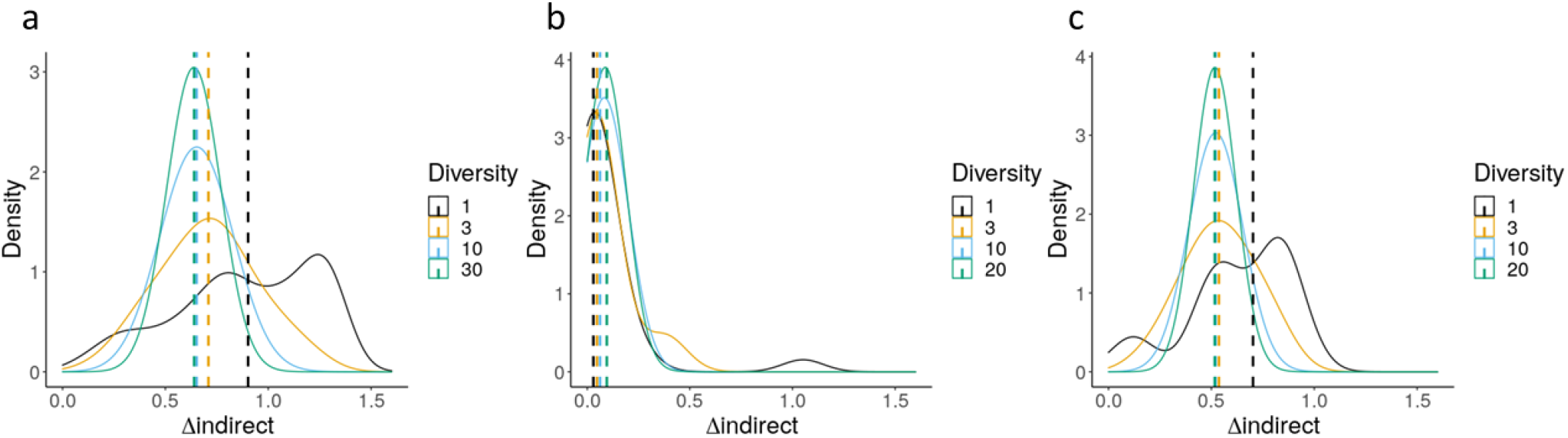
Distribution of indirect effects of warming from 10,000 simulations of a prey species preyed upon by a community of 1-30 predators, with (a) a simulated community of predators (b) a community of ants in California and (c) the same community of ants with *r*_*max*_ = 0.8 for all species. The median of each distribution is displayed by a dashed vertical line.

Increasing the s.d of predator thermal optima increased the effect of diversity on Δ_*indirect*_ up to an s.d. of ~8 (Figure 3c-d), and changing the mean of predator thermal optima increased the effect of diversity of Δ_*indirect*_ when the mean was closer to the prey thermal optimum in terms of survival (17.66 °C; Figure 3a-b). Increasing the breadth of predator thermal niches decreased the effect of diversity on Δ_*indirect*_ (Figure 3e-f).

**Figure 3.**
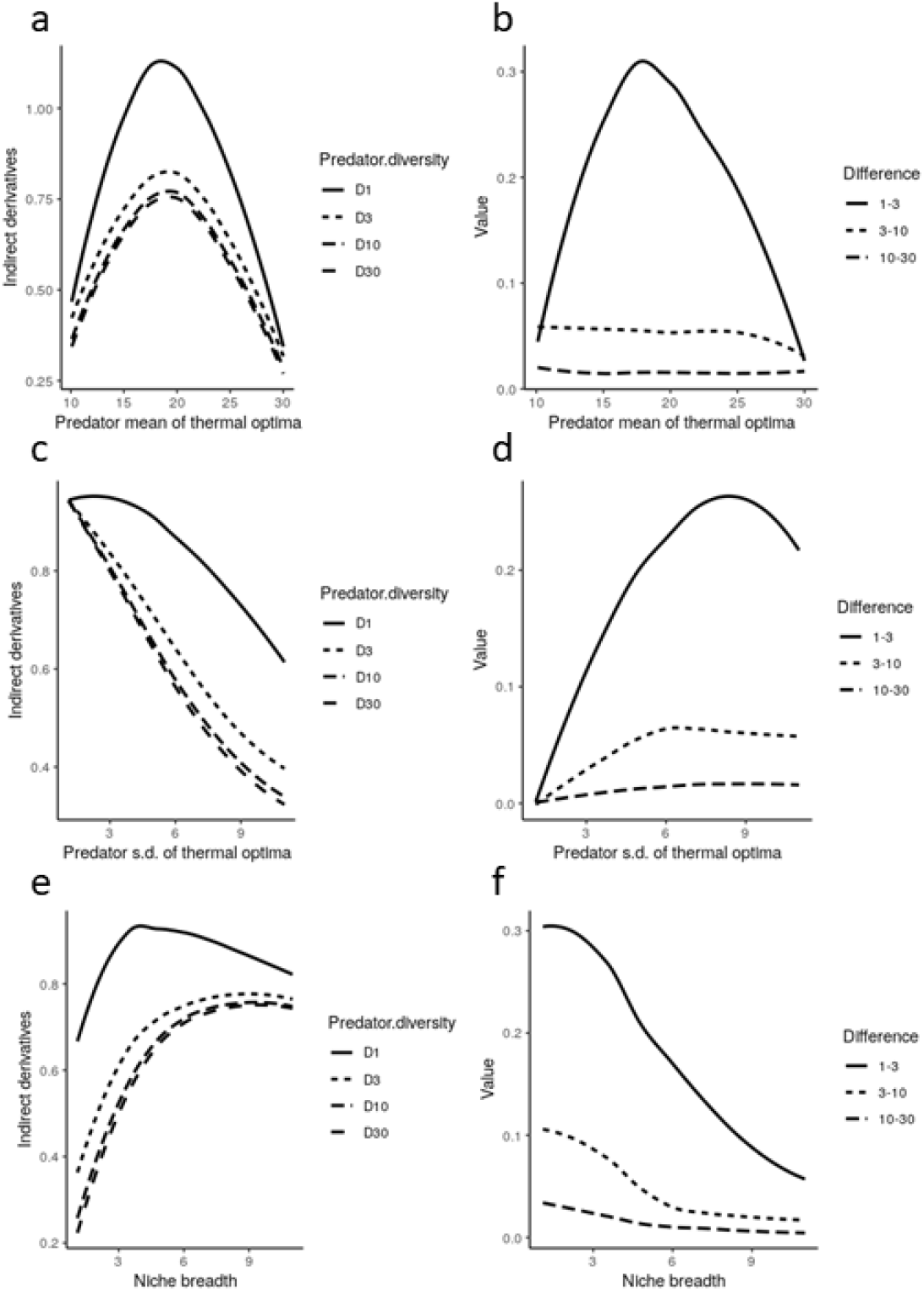
Sensitivity analysis of simulation model, showing (a,c,e) median of Δ_*indirect*_ for communities of 1, 3, 10 and 30 predators, and (b,d,f) differences between 1 & 3, 3 & 10, and 10 and 30 predators. Sensitivity analyses are shown with respect to mean of predator thermal optima (*T*_*opt*_: a,b), standard deviation (s.d.) of predator thermal optima (c,d), and predator thermal niche breath (*σ*_*opt*_: e,f).

## Discussion

We show in our simulations, using simulated and fitted parameter values from a predator-prey community of ants and a caterpillar, that increasing predator diversity may under some circumstances attenuate indirect effects of warming on prey species. The attenuation of the indirect effects of warming we define here as the change in survival probability due to changing interactions with predators (Δ_*indirect*_). We find that this effect is greater when there is sufficient variation in predator thermal optimum and when predators have narrower, non-overlapping niches: that is, when predators have higher thermal niche complementarity. We also find that this effect is greater when the mean of the distribution of predator thermal optima is closer to the thermal optimum of prey survival. Overall, this suggests that higher predator diversity and thermal niche complementarity may buffer indirect effects of warming on a prey species. Our findings also predict that in general, climate warming effects on prey species that are preyed upon by many thermally complementary predator species (i.e., with non-overlapping niches) should primarily be direct as opposed to indirect effects. If generally applicable, this prediction could aid in distinguishing in which circumstances direct effects of warming, which have the potential to be predictable based on species life-history information, are likely to dominate vs. those in which indirect effects of warming would be expected to dominate. This would have the potential for significant value in unravelling some of the complexity of the effects of climate on species when considering trophic interactions (e.g., Post 2013; Dell et al. 2014).

Our modelling approach makes several simplifying assumptions. First, we examine only short-term dynamics, and do not consider reproduction, or long-term outcomes. This is because coexistence of multiple predators in the long term is not simple, since generally whichever predator more efficiently utilizes a prey species will outcompete all the others (Tilman 1982), and studying mechanisms of predator coexistence is beyond the scope of this study. In addition, we do not consider the effects of warming on predator-predator interactions, or varying abundance of different predator species. These are both interesting directions for future research on the effects of warming on trophic interactions; the implications of both however would be to reduce effective predator thermal niche diversity and therefore the attenuation effect on indirect effects of warming that we document. In our simulations, we use the maximal niche diversity for a given predator species richness; that is, an even distribution of all species.

In addition to purely simulated communities of predators, we parameterized our simulation model using empirical data from a community of ants. In this case, rather than using measured attack rates, we use ant activity rates at a given temperature, which is likely to be proportional to attack rate on prey at a given temperature. Due to extreme variation in the rank abundances of ants (see Table S1), we found no effect of diversity on Δ_*indirect*_ with fitted *r*_*max*_ values; however there was an effect of diversity when all *r*_*max*_ values were set to be the same. This effect was smaller than that of simulations not using fitted parameter values because this community of ants consists largely of thermal generalists that have large niche breadths (*σ*_*opt*_) relative to the variation of thermal optima (*T*_*opt*_) and low complementarity (i.e., large niche overlap; Figure S1). The fact the we observed an effect of predator diversity on Δ_*indirect*_ (albeit with adjusted *r*_*max*_ values) even in a predator community that our *a priori* expectation would be for little or no effect of diversity lends credence to the idea that this effect may be common in nature, in communities with similar or higher thermal niche complementarity. In addition, since our measured community includes only species from the same family (Formicidae), it is likely a significant underestimate of the thermal niche diversity in the full community of arthropods at this site, which includes taxa with generally very different thermal niches than ants (e.g., hemipterans, spiders, solifugids, etc.).

In this study we emphasize the concept of the community thermal niche, first developed by Kühsel and Blüthgen (2015), which refers to the collection of the thermal niches of all the individual species in the community. In light of what we have shown in our simulations, we believe that the community thermal niche is a valuable concept to help understand how communities are likely to respond to climate warming, and that a collection of such community datasets would significantly advance our understanding of the ecological effects of warming. There has been a great deal of both empirical (e.g., Brown et al. 2004; Kingsolver and Huey 2008; Kingsolver 2009; Dell et al. 2011) and theoretical work (e.g., Rall et al. 2010; Dell et al. 2014; Uszko et al. 2017) on the effects of temperature on fitness, species interactions, and even simple food webs (Petchey et al. 1999, 2010; Barton and Schmitz 2009). However, the extension of thermal ecology to thermal community ecology should be a priority since individual species and species interactions exist within broader food webs and ecological communities. With the theoretical perspective developed here, some predictions can be made about which aspects of community thermal niche might be important for outcomes with a warming climate.

The range of variation of thermal optima in a community is the primary aspect of the community thermal niche that we expect to mediate effects of warming on a community. As identified by Kühsel and Blüthgen (2015), greater variation in thermal optima in a community represents higher functional diversity, which has been predicted by theory to create greater resilience to environmental change, which they found to be the case for the communities of pollinators that they studied. In the present study, we show further that large variation of thermal optima in a community increases the potential for predator diversity to attenuate indirect effects of warming on prey species. This has the potential to aid in predictions of the effects of warming on predator-prey dynamics: if a community has sufficient variation predator thermal niches, then we can expect that climate will have a greater effect on prey species via direct effects than indirect effects.

The breadth of thermal niches within a community is another aspect of the community thermal niche that we expect to have important implications for the effects of warming on a community. For community resilience, greater thermal niche breadth has a positive effect, since greater breadth of function can only increase resilience (Kühsel and Blüthgen 2015). Kühsel and Blüthgen (2015) define thermal niche complementarity for their purposes as the weighted coefficient of variation of thermal optima. Since we are interested in the degree of change in predation risk with temperature, complementarity here results from a combination of sufficient variation in predator thermal optima as well as narrower predator niche breadth (Figure 3c-f). Narrower thermal niches with sufficiently different optima result in less change in predation risk with temperature because as one predator becomes less active, there is another that is becoming more active, resulting in a consistent predation risk despite changing temperature. In this way, high complementarity combined with greater species diversity leads to much weaker indirect effects of warming.

From here, we can ask in which communities do we expect to see narrower thermal niches, greater variation in thermal optima, and greater diversity of predators, and therefore weaker indirect effects of warming? Generally, we expect to see broader thermal niches (Janzen 1967; Deutsch et al. 2008) and lower diversity (Mittelbach et al. 2007) of ectotherms the farther we are from the tropics. If ectotherms in temperate environments also have higher variation in thermal optima than in the tropics, it is more difficult to make a prediction whether the tropics or the temperate zone will be more dominated by direct or indirect warming effects on predator-prey interactions. This is because greater variability of thermal optima in temperate ectotherm communities would be counteracted by greater niche breadth, just as tropical communities would have their narrower niche breadth offset by a lower variation in thermal optima, in terms of the attenuation effect of predator diversity on indirect effects of warming. In this scenario, whether temperate and tropical ectotherm predator communities have similar or different thermal niche complementarity might depend on if the relative magnitude in the increase of niche breadth and variation in thermal optima with latitude are similar or different enough to result positive, negative, or no change in complementarity with latitude.

We also found from sensitivity analyses that decreasing distance of the thermal optimum of prey survival (in our simulations, 17.66 °C) from the mean of predator thermal optima both leads to greater overall indirect effects of warming and greater attenuation of indirect effects by predator diversity. This means that the thermal optimum of a prey species relative to the community of potential predators has important implications with respect to climate warming effects. If we assume that a species has an invariant thermal niche across its range (which may or may not be the case; Angilletta Jr et al. 2002), and that most species are best adapted to thermal conditions at the climatic mean of their range, then on average a species near its range limit is likely to have a lower thermal optimum than most other species in the community at the warmer edge, and a higher thermal optimum than most other species at the colder range edge. If this is the case, then we can expect much stronger indirect effects of warming at range edges, due to more consistent directional asymmetries and therefore weaker compensating effects of predator diversity. Such effects could contribute to range shifts due to climate warming, through negative indirect effects at warmer range edges, and positive indirect effects at the colder range edge.

## Conclusions

Theoretical and empirical work investigating the effects of climate warming on species interactions has made significant progress in describing how climate warming is expected to affect ectotherm predator-prey pairs (e.g., Dell et al. 2014; Uszko et al. 2017). Little work has extended to examine the effects of warming on multi-species interactions (but see Barton and Schmitz 2009). We propose that the community thermal niche (Kühsel and Blüthgen 2015) is a useful concept and approach to investigate community dynamics under climate warming, based on predictions with regard to community thermal niche traits from our simulation models developed here. Specifically, we predict that species preyed upon by many thermal complementary predators will experience weaker indirect effects of warming than those that are preyed on by fewer or less complementary predators.

Predictions that can be derived from traits of species composing a community are essential if we are to decompose some of the complexity of climate warming effects on communities that are mediated through biotic interactions (cf. Tylianakis et al. 2008; Post 2013; Dell et al. 2014).

---

**Table S1.**
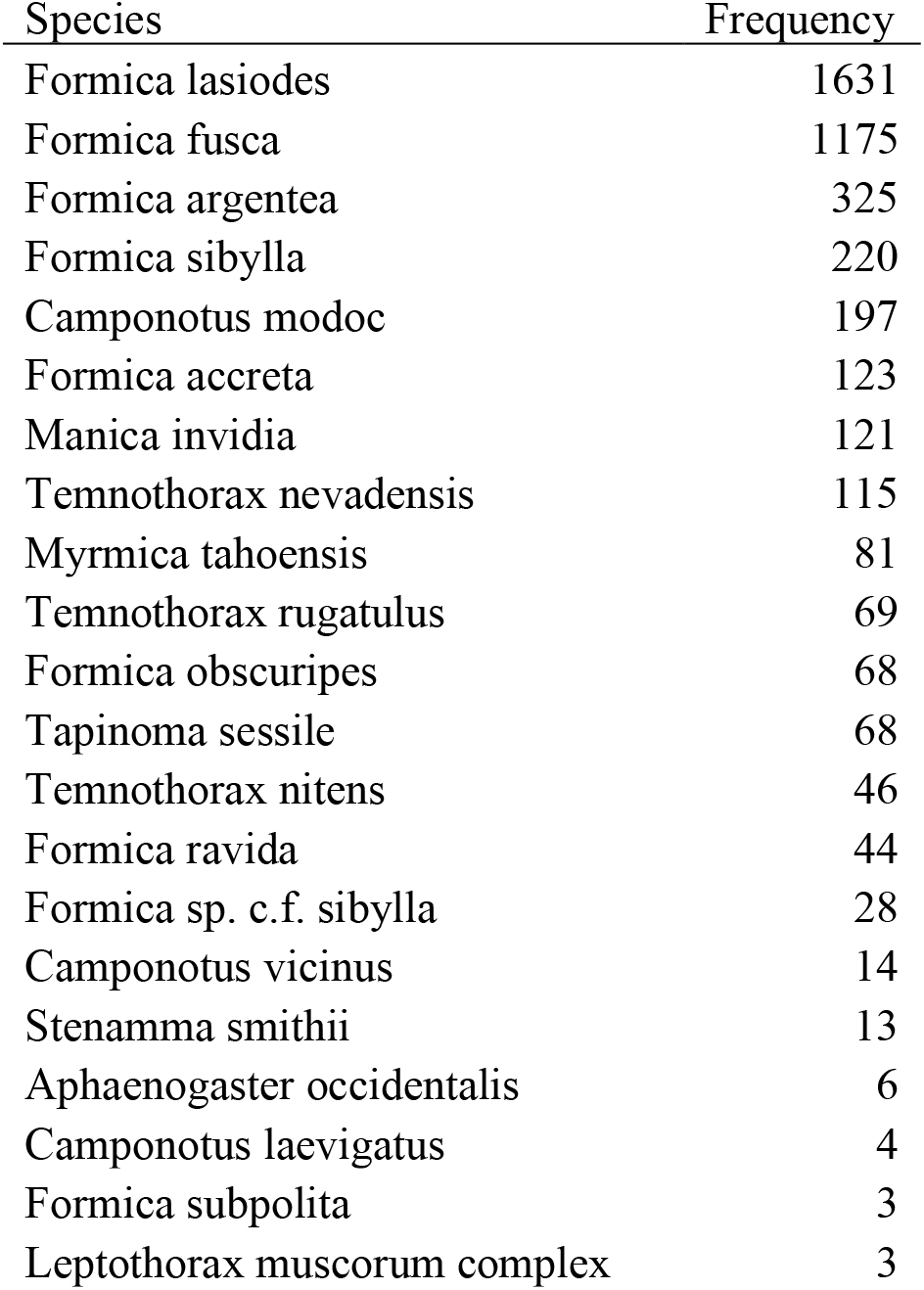
Ranked frequency of ant species in collections, across all times and sampling locations.

**Figure S1.**
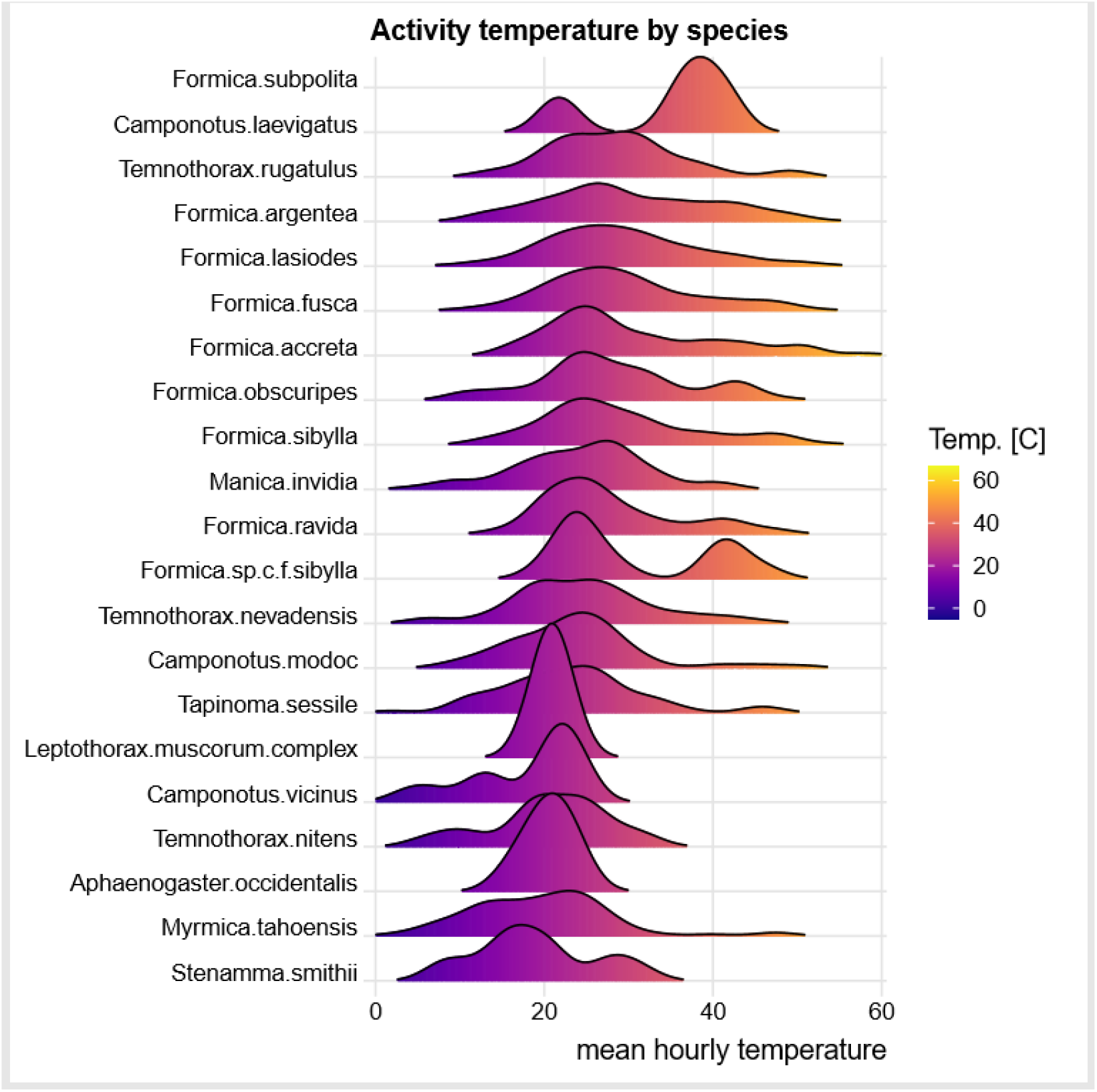
Density plot of ant activity by temperature and species.

